# Social information use promotes breeding synchrony but constrains adaptation to rapid environmental changes

**DOI:** 10.64898/2026.11.20.767538

**Authors:** Jia Zheng, Bram Kuijper, Lucy Aplin, Ben Sheldon, Damien Farine

## Abstract

Animals use social information to make decisions, yet how such processes influence predictions in reproductive timing and population adaptation under environmental change remains unclear. While social information can improve decision accuracy through information pooling, spatiotemporal heterogeneity in conditions and patterns of environmental changes creates noise that potentially limits the value of social information use among breeding individuals. Here, we established an evolutionary individual-based model to examine how social and private sensing interact to shape breeding phenology in species distributed across territories with peaked resources and facing annual fluctuations and rapid climate change. We show that when individuals incorporate social information from neighbours, behavioural responses become increasingly synchronised through positive feedback in decision-making. This emergent synchrony helps populations to adapt faster and can increase reproductive output under stable environmental conditions relative to relying only on private information. However, under directional climate change, the same mechanism reduces population responsiveness by constraining variation in individual responses, leading to systematic lag in tracking shifting environmental optima and reduced population reproductive output. Our results demonstrate that social information use can produce a general trade-off at the population level between synchronisation and adaptive flexibility across environmental contexts. Social interaction enhances population performance when environments are either consistent or changing drastically across both spatial and temporal scales. Rather than uniformly improving performance, social interaction may create conditions under which collective behaviour becomes maladaptive when environments change rapidly.

## Introduction

Novel environmental fluctuations introduce ecological disruptions into animal decision-making^1,2^, requiring them to adapt at rates that are at the limits of animal responsiveness^3,4^. For example, many species adjust their decisions about when to reproduce to reflect changes in phenology^5,6^. Aligning the ephemeral resource peak with the period of high nutritional demands in offspring can substantially enhance reproductive success^7,8^. Global climate change advances spring phenology and generates unpredictable year-to-year temperature fluctuations, placing strong pressure on parents to anticipate the phenology peak before breeding begins^9–11^. A series of phenological cues, such as shifts in photoperiod and ambient temperature, plant budding, and emergence of cocoons, are thought to signal the forthcoming peak^12,13^. Many animals appear to have evolved means of sensing such subtle cues. For instance, wild birds are frequently observed synchronising egg-laying within a few weeks to successfully capture the food peak that comes much later, in the nestling phase^14–16^. Yet, the mechanisms underlying this synchrony in breeding phenology remain a perplexing puzzle for biologists, because the relevant ecological cues can vary at the scale of individual experience^17^. One explanation could be that individuals leverage the landscape of information produced by their conspecifics.

Social interaction is related to collective breeding actions^18,19^ and can shape rapid changes in behaviour^20–22^. Many animals use information produced by others (i.e., social information) when making decisions. Aggregating information from multiple individuals can lead to ‘the wisdom of the crowd’^23^ by averaging noisy individual-level estimates (also known as information pooling). When social information use changes individuals’ behaviours, it can produce highly synchronised collective actions that may improve decisions for better survival or breeding^19,24–27^. For example, changes in movement by some individuals trigger changes in movements by followers, ultimately producing emergent patterns that allow groups to make adaptive decisions^24,28,29^. The combination of social information use and social interactions producing emergent outcomes, so-called collective intelligence, is generally considered an advantage of group-living.

Despite breeding independently in exclusive territories, individuals can acquire information about the actions of neighbours both indirectly (e.g. by eavesdropping on their territorial songs^30,31^) or directly (e.g. by visiting their nests^6,18,32^). Such information allows individuals to learn the breeding status of others, which can then trigger reproductive adjustments. These adjustments can subsequently affect their neighbours, potentially reinforcing actions and spreading them across neighbourhoods—marking a process whereby social information use drives population-level processes. Yet, how such interactions influence reproductive decisions and population breeding outcomes has received little attention^33^. This is likely because any common reproductive interest is thought to be minimal relative to the pronounced territoriality in the first place.

Social information is notable for the fact that even a small change in one individual’s behaviour can trigger a cascade of behavioural changes across entire groups or populations^34^. However, unlike in group-living species, where all individuals are currently experiencing the same space, a limitation to social information use in a territorial context is that environmental conditions are typically noisy and highly variable over a landscape. For example, phenological timing experienced by great tits can vary from one territory to the next^35^. Eavesdropping on neighbours might therefore provide conflicting reproductive information. Using this kind of noisy ‘local’ information transmitted among neighbours may mislead individual decisions and ultimately penalise reproductive success^35,36^.

Ongoing climate change adds another layer of challenge for animals to respond to. It creates generally advanced but less predictable phenology that has ‘global’ effects on all individuals. How the interactions between global directional changes, year-to-year variability^36^ and local heterogeneity affect the benefits of social information use in adaptive reproductive adaptation remains unclear. Directional phenological shifts may complicate the impact of social information use, as information feedback among social members can either accelerate the rates of change or keep the population from making changes. In annually fluctuating environments, social interaction may amplify responses to an exceptionally early year, producing population-level changes that become maladaptive in subsequent years. Alternatively, social interaction may promote convergence in individual decisions, preventing extreme independent predictions across the population. Thus, it is important to systematically explore the potential contribution and consequences of social information driving animals to match phenological shifts across environmental contexts.

In this study, we employ an evolutionary individual-based model to explore the mechanisms that shape breeding phenology under layers of environmental context (i.e., spatial variations, year-to-year fluctuations and climatic advance). Ecological phenology in this model is depicted by a peaked resource curve, which individuals sense through cues informing about food abundance over time^37–39^. We consider the performance of two sensing modes: ‘private sensing’ and ‘social sensing’ (Fig. 1B). In private sensing mode, individuals only rely on their daily perceived ecological cues (current resource abundance) to decide whether to initiate breeding or not. In ‘social sensing’ mode, individuals make a daily decision to initiate breeding based on both their privately perceived ecological cues and social cues from the neighbourhood. Social cues reflect breeding states of neighbours, specifically the number of neighbours that have begun to breed. The decision to initiate breeding in relation to available cues is governed by an evolving reproductive strategy, which is subject to mutation and genetic drift. To investigate how environmental contexts shape evolutionary reaction norms and the corresponding breeding phenology under ‘private sensing’ versus ‘social sensing’, we consider four environmental scenarios (Fig. 1B): first, environmental scenarios where all individuals experience a homogeneous environment with either a steady phenological condition (scenario i) or a fluctuating condition (scenario ii) over seasons in an evolutionary timescale; second, individuals are distributed across a heterogeneous spatial landscape that consists of breeding territories with local variations in phenological timing that either stays stable across seasons (scenario iii), or fluctuates across seasons (scenario iv). Finally, in all the environmental scenarios, we simulate how the different evolved populations adapt to rapid climate change. We do so by introducing a short period of directional phenological change after individuals have adapted to the original environment for a sufficiently long period, by annually advancing the resource peaks across territories.

**Figure 1.**
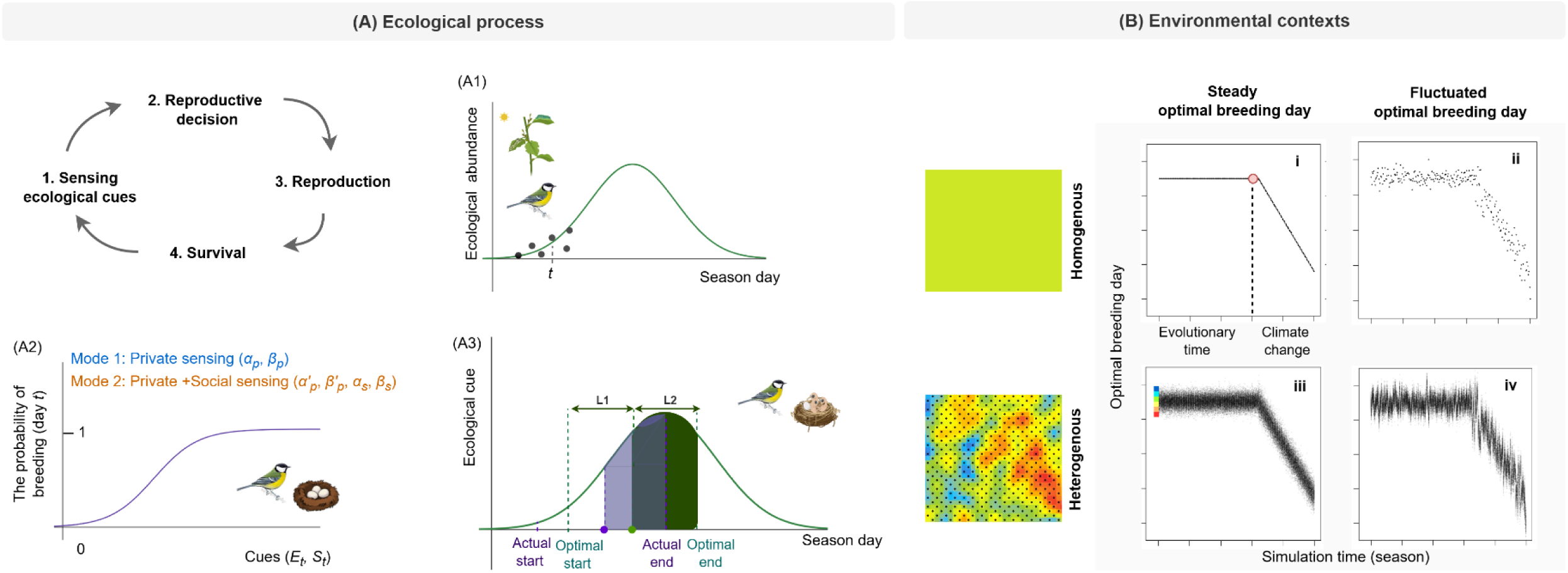
Schematic diagram of model structures and simulation scenarios of landscape and climate conditions. (A) Reproductive process of the model. The breeding season consists of four stages: sensing ecological cues, making reproductive decisions, followed by reproduction and survival. A1--a season begins with individuals sensing daily ecological conditions (reflected by dark dots) to predict an ecological cue. A2--an example of an evolved reaction norm that reflects how individuals make a breeding decision on a daily basis (start breeding or not), either only via ecological cues they perceive and their own reproductive reaction norm (‘private sensing’) or simultaneously consider their neighbours’ breeding status (‘social sensing’). A3--overlapping areas of the provisioning stage to the resource peak determine reproductive success. The length of L1 and L2 indicates the first-half (including the egg-laying and the incubation phases) and second-half (the provisioning phase) in a breeding cycle, respectively. The average reproductive success of a brood is calculated by the size of the shade covered by the actual second-half breeding period (in purple) divided by that of the period centred on the optimal breeding time (in green; this interval is marked as optimal as it maximises the amount of resources). Juvenile and adult survivors will enter the next breeding season. (B) Populations experience a stable climate period for 300,000 seasons (indicated by the pink-dot turning point) and then a rapid climate change period for 150 seasons. We simulated four environmental scenarios: (i) a homogeneous habitat with a consistent phenological peak time (day 20) over seasons; (ii) a homogeneous habitat with fluctuating phenological peak time over seasons. The resource peak time that varies across territories in a heterogeneous landscape. The gradient colour bar from red to blue (on the top-left of B iii) presents the corresponding resource peak times that vary from early to late across territories, and the black triangles on the heterogeneous map indicate individual distributions. (iii) phenological peak times across territories vary across the heterogeneous landscape and stay steady over seasons; (iv) local phenological peak times vary across the heterogeneous landscape and fluctuate over seasons.

## Results

### Private sensing can drive the emergence of a breeding peak

Our private sensing model exemplifies an avian population in which females independently decide the time to initiate breeding (e.g. laying the first egg). In the model structure shown in Fig. 1, ecological abundance follows a seasonal peak, from which individuals perceive daily ecological cues *E_t_* (Fig. 1A1). Ideally, individuals align the nestling development period (the time interval *L*_2_) with the peak resource period. Each individual inherits a condition-dependent reproductive strategy (a behavioural reaction norm, Fig. 1A2) that determines the probability of breeding initiation in response to the local information perceived each day. This reaction norm is encoded by a set of heritable parameters transmitted from parents to offspring, subject to rare mutations (see Methods).

Reaction norms in the private sensing scenario are shaped by two evolutionary traits: *α_p_* is a threshold trait such that an ecological cue value of *E_t_* = *α_p_* yields a probability of initiating breeding of 1/2 at time *t*. The second evolving trait, *β_p_*, is the slope of the reaction norm (initial value *β_p_*__0_ = 0.0), reflecting how the probability of breeding initiation changes with increasing values of the environmental cue *E_t_*. For the sake of tractability, we assume egg laying commences immediately upon the decision to breed, and that breeding cycle length is fixed across all individuals, encompassing egg-laying, incubation (Fig. 1A3, time interval *L*_1_), and provisioning (time interval *L*_2_) stages. Reproductive success is determined by the ratio of the area under the ecological curve during the actual provisioning stage to that during the optimal provisioning stage (the size of the purple shade to the green shade in Fig. 1A3). Individuals who survive the previous season and juveniles who newly join at random will form a fixed-size population and breed in the new season.

In the scenario i (Fig. 1B), we simulate a homogeneous and steady environment. We find that the probability of breeding initiation increases with the perceived value of ecological cues, as positive responsiveness in reproductive strategy evolves monomorphically at the population level (*β_p_* > 0, Fig. 2A, mean± sd: *β_p_* = 9.56±0.78). A breeding phenology peak readily emerges when individuals rely only on private sensing of ecological cues to make breeding decisions (Fig. 2D). Under this strategy, the breeding phenology peak consistently lags behind the optimal breeding time (day 20) across simulation replicates, with reproductive peak on average (mean of the mode ± sd) at 26.50 ± 0.37 days. Sensitivity analysis shows that populations approach the optimal breeding time more closely when seasons are implemented with a broader resource peak (Supplement Fig. 1), suggesting that individuals are better at detecting and responding to a slower rate of resource dynamics over time.

**Figure 2.**
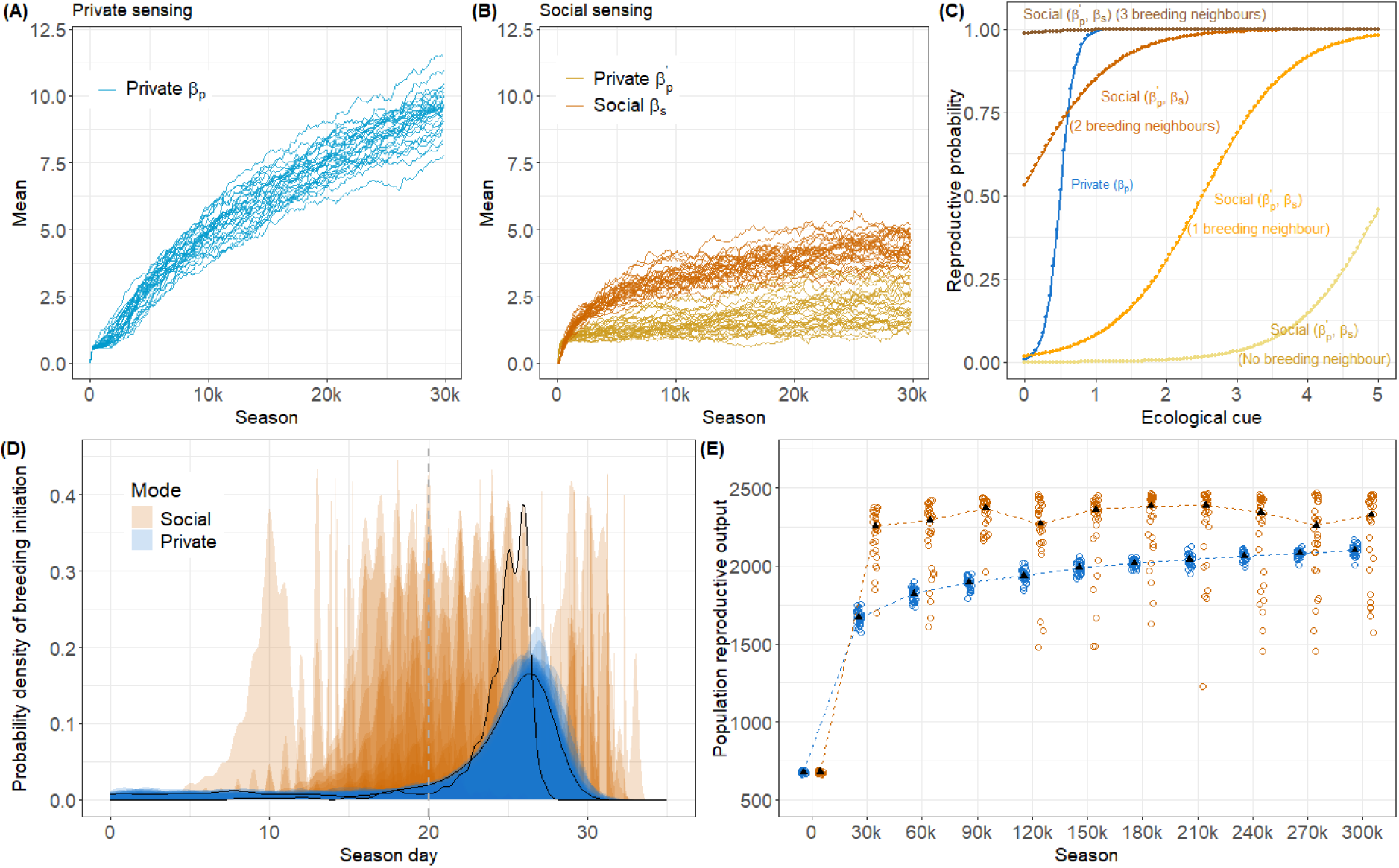
Simulation outcomes of private and social sensing modes over time with annually stable phenological timing (scenario i, before climate change). Both simulation scenarios were run with 30 replicates for 300,000 seasons, respectively. Evolutionary trajectories of the heritable parameters (A) private *β_p_* in private sensing and (B) private 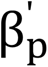 and social *β*_s_ in social sensing. The solid lines indicate the evolutionary trajectories of the population average value of alleles that determined the slopes of reaction norms across simulation replicates. (C) The illustration of reproductive reaction norms evolved from private sensing (blue) and social sensing (yellow-to-brown series) modes using a random simulation outcome. Private sensing norm is shaped by alleles *α_p_* and *β_p_* and social sensing norm is shaped by alleles 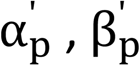, *α*_s_ and *β*_s_. The number of neighbours who are in breeding status increases in the order of yellow, orange, brick and brown curves, showing that individuals rely less on ecological cues to make reproductive decisions when more neighbours begin to breed. (D) Breeding phenology of the last simulation season under private sensing (in blue) and social sensing (in orange) modes of all simulation replicates. The grey dashed line presents the optimal breeding time (Day 20). One pair of breeding phenology emerged under the two sensing modes is marked up to show the brief difference. (E) Population reproductive output during the evolutionary timescale. The hollow circles show outcomes from all simulations, black triangles indicate the median value over simulations, and the dashed lines indicate the trend of output changes over simulation time.

### Social information use improves population adaptation and facilitates breeding synchrony

In the social sensing scenario, individuals can receive additional social information from their neighbours. They detect the number of neighbours that have initiated breeding as a social cue *S_t_*. The probability to initiate breeding is now sensitive to both private and social cues. Sensitivity to the ecological cue is given by evolving threshold 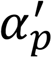 and slope values 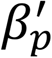, while sensitivity to the social cue is given by evolving threshold *α_s_* and slope values *β_s_* in a sigmoidal function (See Fig. 1A2, more details in the Methods section).

Social sensing likewise evolves so that the probability to initiate breeding is positively affected by social cues, so that individuals are more likely to start breeding once more neighbours have started breeding (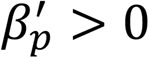 and *β_s_* > 0, Fig. 2A and B). Although both social and environmental cues vary across the same scale, individuals are more responsive to social cues (mean± sd: 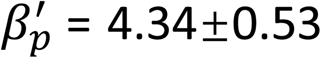) than to ecological cues (mean± sd: *β_s_* = 2.25±0.76, Fig. 2B). This indicates that they rely more on social sensing to decide on breeding initiation. Notably, individuals in the social sensing scenario are much less responsive to ecological cues than in the private sensing scenario (Fig. 2A, mean± sd: 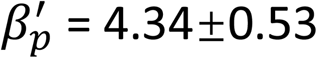 vs *β_p_* = 9.56±0.78). This is directly reflected by the substantially lower probability of breeding initiation when ‘no neighbour’ has initiated breeding (Fig. 2C). However, as more neighbours initiate breeding, individuals’ breeding propensity soars up, such that breeding is initiated immediately once three of the neighbours have begun breeding, independent of ecological cues (Fig. 2C, ‘3 breeding neighbours’).

Incorporating social interaction substantially enhances breeding synchrony. Individuals’ peak timing of breeding initiation moves more closely to the optimal breeding time of day 20, compared to the ‘private sensing’ scenario (75% of simulations, mean breeding phenology modes ± sd: 22.59 ± 5.54 days). Notably, social interactions also introduce more variability in the peak of breeding onset across simulations (Supplement Fig. 5). Social information use also accelerates population adaptation, leading to a faster increase towards a higher reproductive output than in populations that only use private sensing, although with greater variation (Fig. 2E; median across simulations: private = 2101, social = 2326). However, this benefit of social sensing on reproductive output disappears when increasing the resource peak fluctuation across seasons (Fig. 1B, scenario ii, Supplement Fig. 4A3). Increasing the number of neighbours from which individuals acquire social information produces more synchronised breeding phenology, but does not alter population output (Supplement Fig. 3).

### Landscape heterogeneity differs in the distribution of breeding time between sensing modes

We introduce spatial heterogeneity within landscapes (Fig. 1B, scenario iii) at a scale that resembles empirical studies^35^. Individuals who survive the previous season will breed at their original territory in the new season, and newly joining juveniles are allocated to vacant territories at random to breed. Here, the relative timing of the ecological peak is now affected by between-territory noise and being different among neighbouring territories (more details see Methods).

Our results show that the distribution of prediction mismatch (estimated as the difference between the actual and the optimal breeding time) across territories is different between private and social sensing. In purely private-sensing populations, individuals initiate breeding according to the phenological condition of their territory, breeding early where resources peak early and late where they peak late (Supplementary Fig. 5, top row). As a result, the degree of mismatch is independent of the locally optimal breeding time (Fig. 3A). Contrastingly, populations inhabiting the same landscapes but using social sensing tend to show no correlation between actual breeding time and optimal breeding time (Supplement Fig. 5, bottom row). Individuals with an early optimal breeding time tend to breed later than their optima due to few or no neighbours have yet started breeding, whereas individuals with a late optimal breeding time tend to breed earlier, because conformist social effects lead they to follow early-breeding neighbours, which reduces their lag relative to the breeding optima. This produces a negative correlation between breeding mismatch and optimal breeding time. Notably, social effects can over-correct the lag, causing earlier breeding than the optimal time (Fig. 3A).

**Figure 3.**
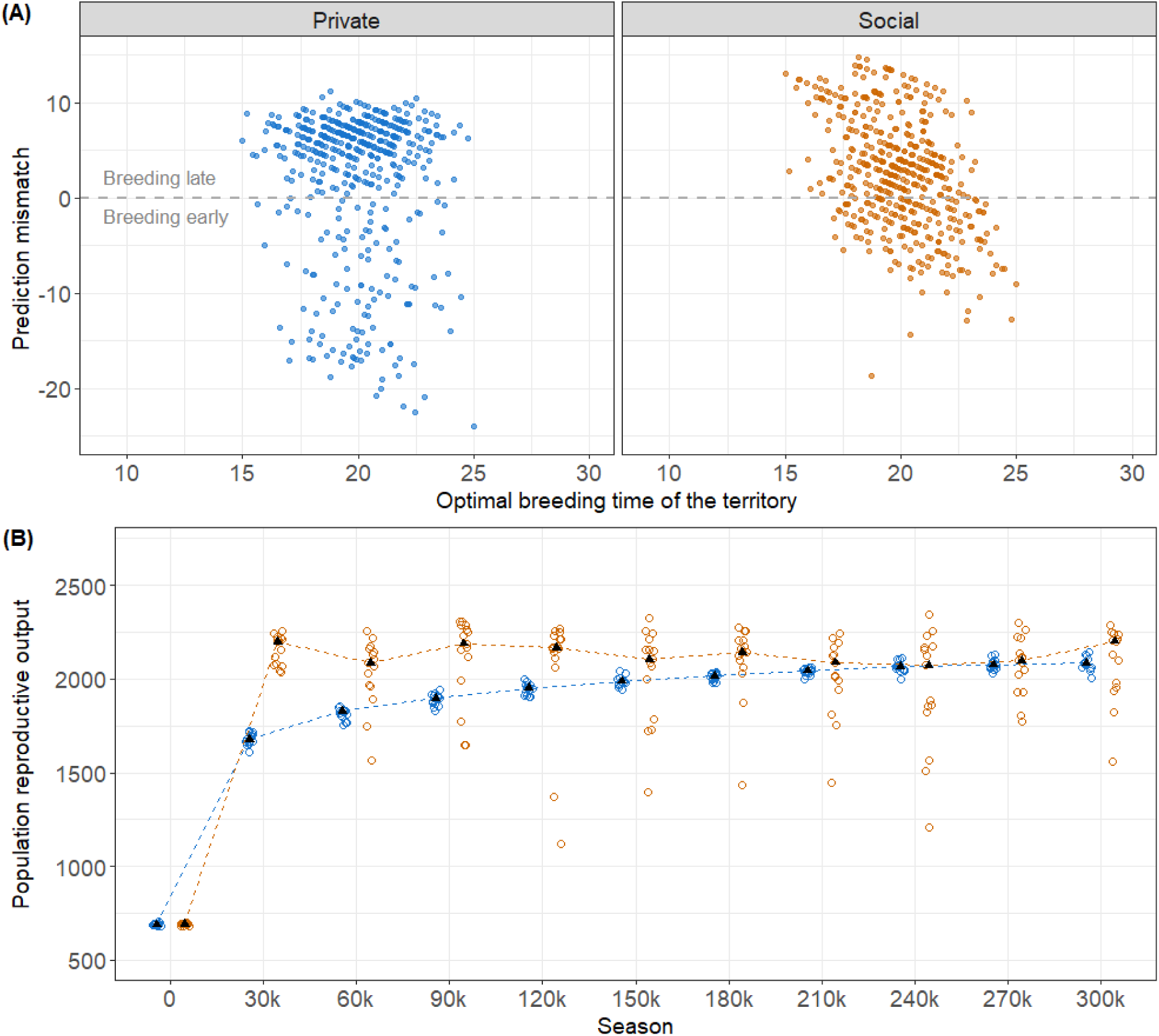
Different distributions in breeding mismatchs but similar population reproductive output under private and social sensing in a stable heterogeneous habitat (scenario iii). Simulation scenarios of private sensing (in blue) and social sensing (in orange) were run for 50 replicates each. (A) Individual prediction mismatch across territories. The result is shown from one random simulation to illustrate the pattern of breeding phenology. Prediction mismatch is calculated by the actual breeding time minus the optimal breeding time in a territory (the grey dashed line). It is consistent in private sensing, but directionally changes in social sensing across territories (see Supplementary Figure 6 for more simulation replicates). (B) Population reproductive output under the two sensing modes converges to similar levels at the simulation end. The hollow circles show the population reproductive output resulting from simulations, the black triangles present the median value across simulations, and the dashed lines indicate the changing trend in population output.

When comparing these two types of populations, we find that the private sensing populations show more outliers towards too early breeding, whereas breeding times are more clustered in social sensing populations (Fig. 3A, Supplement Fig. 5), but show more variable distributions and reproductive output over simulations (Fig. 3B and Supplement Fig. 6). Populations using social sensing obtain similar reproductive output as populations that only use private sensing in heterogeneous environments (Fig. 3B, median: private 2096 vs social 2123). Notably, when increasing year-to-year fluctuations to ecological resources (scenario iv), social sensing again benefits populations by consistently achieving a high reproductive output (Fig. 1B, Supplement Fig. 4B).

### Social information use is not adaptive and negates population performance under climate change

We introduce a rapid phenological advancement of 0.1 day per season for 150 seasons, applied at the end of the simulation (i.e. after season 300K) in both homogeneous and heterogeneous landscapes. We include climate change in both sensing scenarios, with and without annual phenological fluctuations (scenarios i, ii, iii, iv). We find that, surprisingly, private sensing populations adapt better during climate change than social sensing populations, regardless of environmental heterogeneity or annual fluctuations. Private sensing populations track the resource changes by steadily advancing their breeding phenology during climate change in a homogeneous environment (Fig. 4A in blue), showing increasingly clustered prediction mismatch around the optimum (i.e., better breeding timing) across heterogeneous territories (Fig. 4B in blue). This results in a stable and slightly increased reproductive output at the population level (Fig. 5A and B). In contrast, social sensing populations perform much worse in tracking the optimal breeding time during climate change, showing highly variable breeding phenological peaks (Fig. 4A in orange) and consistently large breeding time mismatch across territories to the optimal time (Fig. 4B in orange), resulting in substantial declines in population reproductive output as climate advances (Fig. 5A and B). Faster climate change deteriorates the adaptation of social sensing populations, but does not affect that of private sensing populations (Supplement Fig. 7, A2). In both the social and private sensing scenarios, year-to-year fluctuations destabilise population reproductive output (showing larger variations) in a homogeneous landscape (Fig. 5B), but do not affect the reproductive output of populations distributed in a heterogeneous landscape (Fig. 5D). The rate of climate change relative to between-year fluctuations has consequences for adaptation and reproductive output. Differences between the two sensing modes are reduced when annual fluctuations increase (Supplement Fig. 7, B1 and B2).

**Figure 4.**
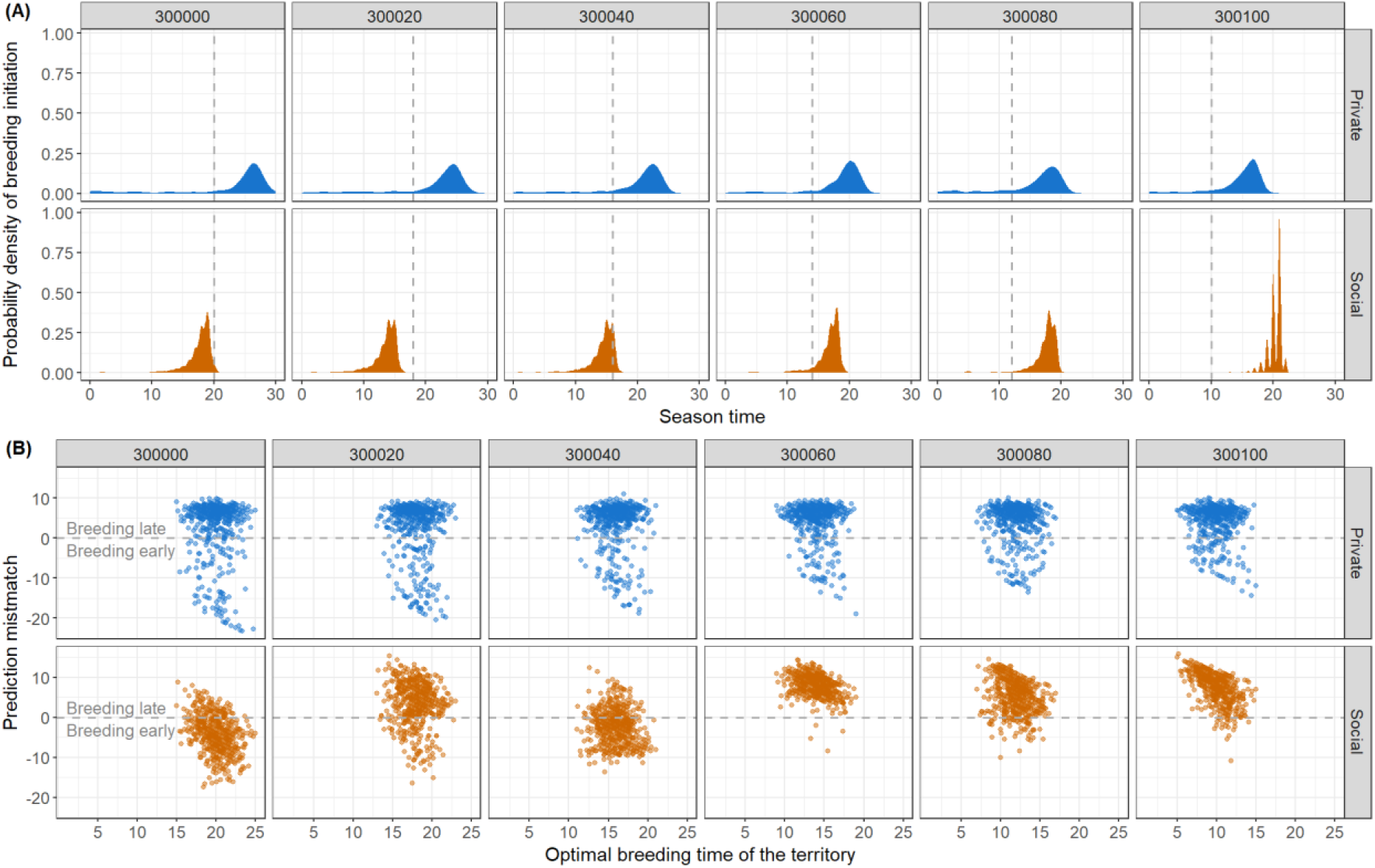
Illustration of the distributions of breeding time during climate change in homogeneous and heterogeneous simulations (scenarios i and iii). Simulation outcomes of private sensing are in blue, and those of social sensing are in orange. (A) Breeding phenology during climate change in a non-spatial model. The grey vertical dashed line shows the optimal breeding time during climate change; (B) Prediction mismatch in breeding time of individuals distributed across territories with varied optimal breeding times during climate change. The grey horizontal dashed lines (y = 0) represent an ideal prediction in line with the territorial optimal breeding time. For results of more simulation replicates of the distribution of actual breeding time, please see Supplement Fig. 8 and 9.

**Figure 5.**
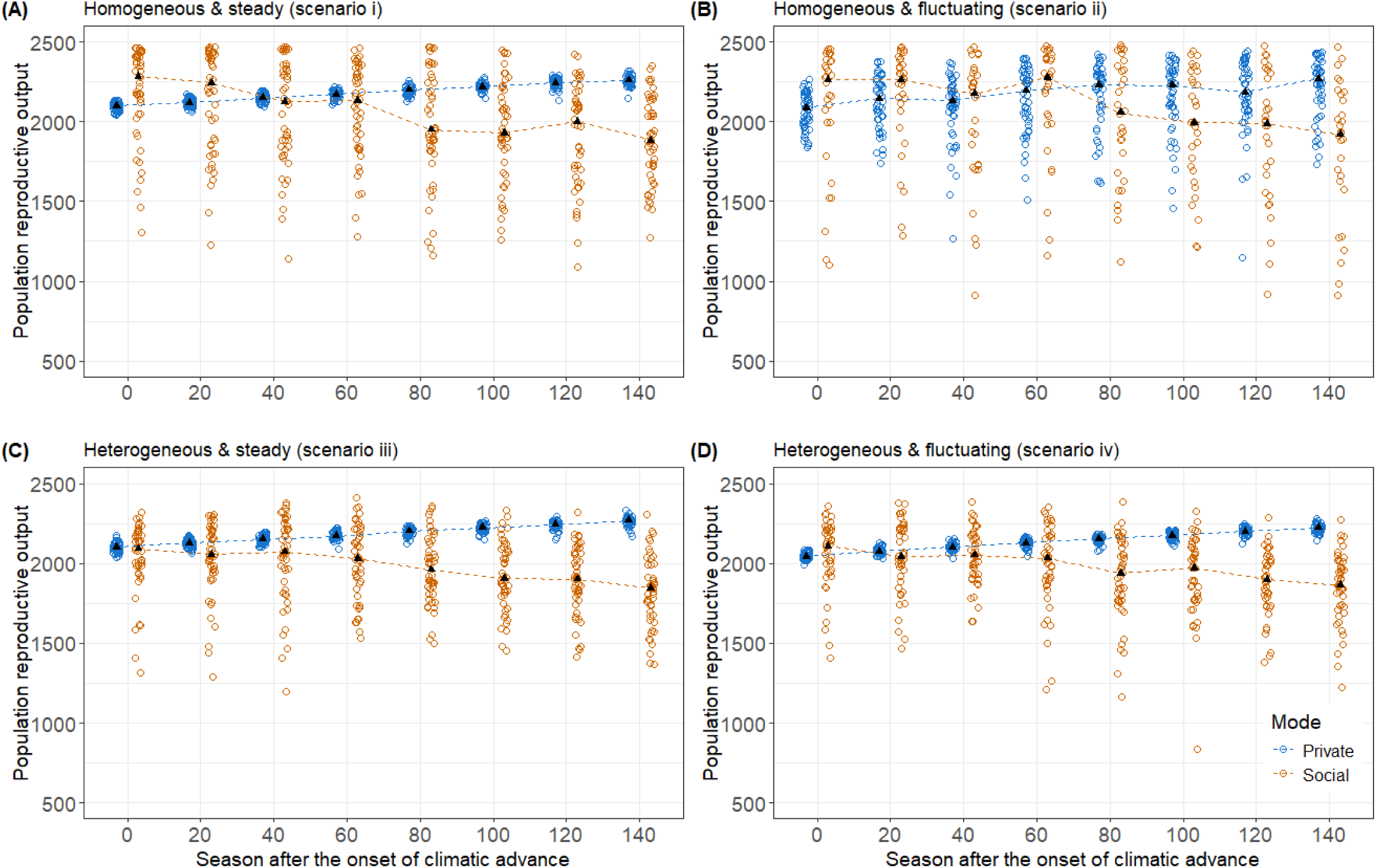
Population-level reproductive output during rapid climate change under steady and fluctuating environments in homogeneous and heterogeneous landscapes. Simulation replicates were run 50 times each under the private (in blue) and social (in orange) sensing modes, respectively. Population reproductive outputs under different simulation scenarios were presented every 10 seasons during climate change in four environmental scenarios: (A) scenario i: homogeneous and steady; (B) scenario ii: homogeneous and fluctuating; (C) scenario iii: heterogeneous and steady; and (D) scenario iv: heterogeneous and fluctuating. The hollow circles show the population reproductive output resulting from simulations, the black triangles present the median value across simulations, and the dashed lines indicate the changing trend in population output.

## Discussion

Animals must make breeding decisions in response to resource dynamics and environmental change^13,15^. These decisions are often driven not only by their own estimations of environmental conditions, but also by socially acquired information^21,40,41^. Our study presents the first theoretical model considering the evolution of ‘private sensing’ and ‘social sensing’ in shaping population breeding phenology across complex environmental contexts. We found that social information use substantially increases breeding synchrony, enables populations to rapidly converge on a timing strategy that leads to high reproductive output, and reduces the systematic lag in breeding timing relative to the optimal resource peak. At the same time, the conformist effects of social interactions drive substantial variation in the timing of breeding peaks across simulation replicates, reflecting the amplifying role of early breeders on subsequent population-level responses. In the presence of spatial heterogeneity, social interaction creates ‘information noise’ transmitted among neighbours. This noise reduces the advantage of social sensing under temporally stable conditions, while its benefit on reproduction is restored in populations experiencing larger annual fluctuations. Across environmental configurations, the effect of social information use on population reproductive output, and the variation in this output, depend critically on the interaction of spatial heterogeneity and annual fluctuations. Strikingly, during rapid climate change, social sensing populations adapt substantially less effectively than private sensing populations and uniformly show decreasing reproductive output across all the environmental scenarios.

Breeding phenology peaks are commonly observed in wild birds^6,16,19,42^. Despite considerable efforts to analyse breeding phenological adaptations through field experiments and meta-analyses ^8,48,49^, the evolutionary processes underlying the formation of breeding peaks and population responsiveness to environmental change remain largely inaccessible to empirical investigation alone. Evidence shows that phenological timing traits are heritable and shaped by selective pressures^43–46^, enabling territorial wild passerines to plastically adjust their egg-laying dates in response to local phenological changes over decades^5^. Our simulation model captures this eco-evolutionary dynamic by allowing individuals to make breeding decisions via an explicit evolutionary reaction norm, governed either by temporally varying ecological cues alone or alongside social cues from neighbours. This framework bridges breeding decisions at the behavioural timescale with the heritable sensing strategies over evolutionary time, revealing how their interplay shapes the population-level breeding phenology in nature and their capacity to respond to environmental change.

Our results show that breeding phenology peaks readily form through private sensing, but consistently lags behind the optimal timing given a narrow resource peak. Social information compensates for this lag through a positive feedback mechanism: as early breeders trigger neighbours to breed, conformity spreads across the population, reinforcing reproductive synchrony. This emergent synchrony resembles collective intelligence seen across animal systems. For instance, an ecological model of fish populations similarly found that incorporating social cues triggered more synchronous spawning that better matched empirical breeding phenology^50^. The amplitude of breeding synchrony also increase with the size of social neighbourhoods, consistent with the positive association between synchrony and group sizes observed in colonial bird species^18,51^.

It is controversial whether social interaction benefits individuals and the group^52–54^, as social effects can increase decision similarity at the cost of diversity among private predictions^26,54^. We found that introducing social interaction substantially increases variation in both breeding peak timing and reproductive output across simulations compared to private sensing alone. This variation is largely attributable to the predictive precision of early breeders: when early breeders happen to time poorly, this mismatch is propagated across the population regardless of ecological cues. This produces a long-tailed distribution of population output across replicates, where most simulations perform well but the outliers experience extreme deviations in breeding peak timing driven by poorly-timed early breeders. Nevertheless, across most simulations, the benefits of social sensing in fast adaptation and increasing population reproductive output outweigh the costs of increased variation, reflecting a broader trade-off between synchrony and adaptive flexibility arising from social information use.

Annual fluctuations and spatial heterogeneity reflect two selective pressures that jointly shape the evolution of reproductive reaction norms and breeding phenology. We found that the interaction between spatial heterogeneity and annual fluctuations reveals a broader principle governing the role of social information use: its benefit depends critically on whether the selective pressures imposed by these two environmental layers are directionally aligned. When they are concordant (either both generating stochastic variations--scenario iv or both imposing stable conditions—scenario i), neighbours’ breeding behaviours collectively constitute a more reliable signal. Under universally stochastic conditions, social information aggregates noisy individual estimates and restores a population-level average that tracks the shifting optimum (the “wisdom of crowds” effect), whereas universally stable conditions allow social information to consistently reinforce convergence on a fixed optimum. By contrast, when the two environmental layers are discordant, such as when habitats are spatially heterogeneous but temporally stable across seasons, the selective pressures they impose pull individual strategies in opposing directions. Spatial heterogeneity favours sensitivity to local ecological cues, while temporal stability selects for precise, fixed strategies. In this conflict, social information transmitted among neighbours introduces systematic bias rather than correcting individual estimates, ultimately dampening the benefits of social interaction. This interpretation is consistent with theoretical work showing that optimal cue integration under environmental uncertainty requires congruence between the timescales at which different information sources operate^37^. Our results offer a theoretical lens through which contrasting phenological patterns across species may be interpreted: spatially heterogeneous habitats with phenologically fluctuating seasons impose concordant pressures that enable territorial passerines to fine-tune with local phenology and achieve stable population output^35,56^; in contrast, seabirds breeding in homogeneous colonies while experiencing year-to-year fluctuations in aquatic environments face discordant pressures, consistent with the distinctively unstable breeding phenology and decreased population performance observed in these species^36,57^.

Surprisingly, during the climate change period, which exhibits continuously advancing phenology, social sensing consistently results in population reproductive reduction regardless of environmental context. This type of directional environmental change has been characterised as “red noise” — a series of environmental variations that unfold rapidly and are not entirely independent of prior values — and has been associated with fitness decline and population collapse in nature^58^. Unlike the stochastic or stable conditions under which social sensing proved beneficial, red noise imposes a unidirectional selective pressure that fundamentally conflicts with the conformist dynamics of social information use. The mechanism underlying this maladaptation lies in an “evolutionary trap”. During the prolonged period of environmental stability preceding climate change, reproductive strategies in the population evolved towards social conformity over private prediction (shown by 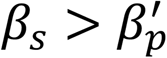). This conformist strategy, while advantageous under stable or concordantly variable conditions, reduces population-level timing variation to track a shifting optimum. When rapid climate change arrives, the positive feedback inherent in social sensing amplifies responses in the wrong direction, preventing the population from tracking the advancing resource peak. By contrast, populations relying solely on private sensing retain greater capacity to track shifted ecological cues 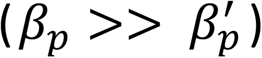, maintaining consistent productivity over time. This dynamic resonates with empirical observations: passerine birds have been found to advance their laying dates in response to warmer springs^49,59^, yet consistently lag behind the shift in resource timing^51^. This is a pattern of maladaptation recognised across species, but its underlying mechanisms remaining unclear^60,61^. Our results offer a new mechanistic explanation: social interactions, while facilitating adaptation under environmental complexity, may represent a short-sighted strategy that trades long-term adaptive flexibility for short-term synchrony gains, ultimately threatening population stability when environments change rapidly and directionally.

Social information use represents a previously overlooked driver of population vulnerability to environmental change. Our model provides the first mechanistic exploration of how social information use shapes breeding phenology across environmental contexts. By bridging behavioural decision-making at the seasonal timescale with the evolution of sensing strategies over evolutionary time, our framework reveals how the interplay between social and private information use generates emergent population-level outcomes that cannot be predicted from individual-level processes alone. Critically, our results demonstrate that the effect of social information use on population performance is not fixed, but depends on multiple layers of environmental contexts and their interactions. We established this model around a fundamental mechanism realised in many avian species that time reproduction to coincide with the resource peak critical for offspring survival. Naturally, breeding phenology in nature is also shaped by additional processes not included here. For example, species-specific life-history and demographic traits drive variation in responses to environmental change^8,11,51^, interactions among sympatric species can mutually influence breeding timing^6,48^, and alternative mechanisms such as predator dilution or early-fledging competition may further complicate phenological dynamics^33,62^. These simplifications are deliberate, as our aim was to isolate the evolutionary consequences of social information use across environmental contexts, rather than to replicate the full complexity of any particular system. Our reproductive reaction norm framework offers a multivariate space that can readily incorporate co-evolutionary traits, species-specific sensing mechanisms, and additional ecological pressures. We encourage future studies to build upon this foundation, as doing so will deepen our understanding of the evolution of breeding phenology and improve our capacity to predict species responses in a rapidly changing world.

## Methods

### Phenology and sensing ecological cues

In this model, a breeding season begins with the emergence of a phenology curve that reflects the dynamics of actual ecological resources, such as the number of buds, the encounter frequency with insect cocoons, and ambient temperature. This phenology curve *R* is normally distributed over an entire breeding season that lasts for 80 days (Fig. A1, eq. [1]). The standard deviation σ of this normalized phenology curve is 10 (see Supplement Fig. 1 with accompanying rationale regarding the values of σ), and it peaks on the day *μ* (= 50) at *R_μ_* (= 8), to ensure the resource distribution has the same data scale as the number of neighbours (see ‘Sensing social cues’ below for more information), allowing one to compare social and environmental reaction norms.

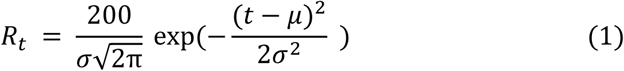

Individuals perceive an ecological cue *E_t_* from a normal distribution that is averaged by the actual resource value *R_t_* and varies with a prediction noise of *R_t_* * (*E*^−0.3*t*^ + 0.1), reflecting the assumption that the absolute value of the prediction noise is relatively larger for a low resource amount, as resource scarcity often coincides with higher predictive uncertainty^37^. Individuals sense ecological cues on a daily basis until they decide to initiate breeding.

### Sensing social cues

In the social sensing scenario, individuals make reproductive decisions by incorporating cues from both ecological and social sensing. Here, each individual has *N* (= 8) neighbours (*N* and the peak of ecological cue *R* have the same scale, to simulate a comparable selection strength on the private and social sensing, for sensitivity analysis on the number of social neighbours, see Supplement Fig. 3). Neighbours are assigned at random and following one rule: If individual A is allocated neighbour B, then B also recognizes A as its neighbour. However, A’s other neighbours (for example, C) may not appear among B’s neighbours. Individuals obtain social cues by counting the number of neighbours who are in the breeding state *S_t_*. They will then use both their perceived ecological cue and social cue to make their breeding decision (see eq. [3] below).

### Breeding decisions

In the private sensing scenario, breeding decision is governed by the daily sensed ecological cue and an inheritable reproductive strategy. We consider this strategy of the form:

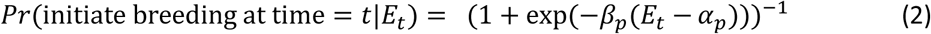

As shown here, an individual’s reproductive probability on a day *t* is determined by its private-sensed ecological cue *E_t_* via a sigmoidally-shaped reproductive strategy. This strategy is a logistic function that ranges between zero and one and is characterised by the two evolvable parameters *α_p_* and *β_p_* carried. Parameter *α_p_* represents the value of the ecological cue that leads to a 50% breeding probability, and *β_p_* is proportional to the steepest ascent of the reproductive function. In our model, the initial value of *β_p_* is zero, representing a neutral and non-responsive state to the environment as the initial condition. The sign of the evolved *β_p_* determines whether reproductive probability increases or decreases with the ecological cues. The initial value of *α_p_* is 5 (adjusting the initial value of *α_p_* appears not to influence the evolutionary outcomes (Supplement Fig. 10). Individuals in our models are haploid and asexual to resemble a female-dominant timing of the egg-laying.

For the social sensing scenario, individuals carry two more parameters *α_s_* and *β_s_* that shape the reproductive strategy together with 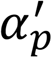 and 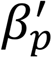 (corresponding to *α_p_* and *β_p_* in the private sensing scenario) in the logistic function reflecting the probability to initiate breeding at time *t*:

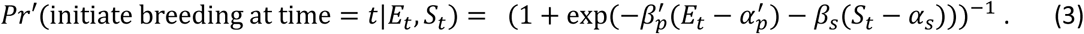

The initial values of the reaction norm slops 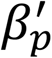 and *β_s_* are both zero. When slopes evolve to any nonzero value, the daily sensed ecological cue *E_t_* and social cue *S_t_* (reflecting the number of neighbours in breeding status; Farine 2022) simultaneously affect the probability to initiate breeding. In both private and social sensing scenarios, an individual’s decision to initiate breeding on day *t* is then drawn from a Bernoulli distribution with probabilities in private and social sensing scenarios given by eqns (2) and (3) respectively.

### Reproduction, survival and strategy inheritance

Females often determine the initiation of egg-laying in birds^32^. For the sake of tractability, we consider a haploid, asexually reproducing population. Reproduction starts once an individual decides to breed on the day *d*. Each clutch has *C* (= 5) eggs and the entire breeding cycle lasts for 40 days, during which the first half cycle (= 20 days) is spent on egg-laying and incubation, and the second half on provisioning offspring. Since resources are intensively demanded by hatchlings, broods whose midpoint of the provisioning stage more closely approaches the phenology peak *d_pEak_*(= 50 in the steady environment) obtain higher reproductive output. This makes the optimal timing of provisioning from *d_pEak_* − 10 to day *d_pEak_* + 10, thereby the optimal breeding starts at *d_pEak_* − 30. ss eproddcctiee roddcctiiiyy aas oftnn umpp-shaped against season time in birds^51^, we calculate reproductive success *W_i_* of individual *i* using the integral of the phenology function *R* during the actual provisioning period, to divide that by the optimal provisioning period,

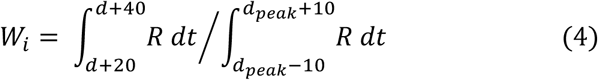

Each individual has one brood at most in a season, so their reproductive output in the season is determined by *W_i_* * *C*. We consider a population with a consistent size of 500 over seasons, to resemble the limited territories in a natural habitat. At the end of the season, all individuals survive to the next season with a given probability of 0.5. Non-surviving adults are removed from the population and replaced by the newly-born offspring, parents of whom are sampled from the distribution of current season’s breeders, in which each breeder is weighed by its reproductive success. Offspring adopt the reproductive strategy encoded by inherited, non-linked haploid alleles (*α_p_*, *β_p_*) or (*α′_p_*, *β′_p_*, *α_s_*, *β_s_*) from their parents. All the alleles are subject to a per-generation mutation probability of *μ* = 0.03, and the mutational step size is drawn from a normal distribution with a mean of zero and a standard deviation of σ = 0.1. This mutation is added to the corresponding parameter value that the offspring inherited. All surviving adults and offspring enter the next season for breeding.

### Establishing a heterogeneous landscape

We later employ the main model in a spatial landscape containing 33×33 grid cells, each representing a potential breeding patch. To produce spatially correlated heterogeneity, we used a multi-scale fractal noise approach based on Perlin noise^63–65^ Specifically, each grid cell value was determined by summing multiple layers (“octaves”) of Perlin noise with progressively decreasing amplitude and increasing frequency. The contribution of each layer was controlled by persistence, which determines how much each layer contributes to the overall landscape, and clumpness, which scales the extent of spatial change between successive layers. The raw noise values were normalised to ensure consistency across replicates and rescaled to a biologically meaningful range^35^. The value in each cell represents phenological peaks spanning 10 units centred around the mean of 50 (corresponding to the phenology peak in the main model). This approach produced spatially structured but smoothly varying landscapes, reflecting environmental gradients in nature.

### Simulation in the spatial landscape

In this simulation scenario, potential territories are distributed every second grid cell in both the x- and y-directions in a regular lattice pattern. At the beginning of each season, adult individuals who survived through the last season will stay at the previous territory, whereas the newcomers entering the season will be randomly assigned to vacant territories in the landscape. We also separately consider the private and the social sensing scenarios in this spatial simulation. In the social sensing scenario, they acquire social cues by inspecting the breeding status of the eight neighbours residing in the surrounding territories. Some individuals located on the edge of landscape grids may have fewer neighbours. Breeding decision, adaptation of reproductive strategy and progression of a breeding season were all implemented accordingly as in the homogeneous model. New seasons begin with shuffling all individuals, who are later randomly allocated to breeding territories.

### Simulating process, climate change and fluctuation

Overall, we ran hundreds of simulations for all four model analyses: homogeneous private model, homogeneous social model, spatial private model, and spatial social model. In all scenarios, we first implemented the models at an evolutionary timescale (i.e. for 300,000 seasons) to explore the evolution of reproductive strategies. This course of time is sufficient for the emergence of a stable reproductive strategy. Climate change is involved subsequently and persists for 100 years, during which the phenology peak advances by 0.1 day each year. In addition, we simulated scenarios with fluctuating climate conditions in all four model analyses by adding noise to the phenology curve during the entire evolutionary and climate-changing stage. Annual fluctuations during the evolutionary period (first 300,000 seasons) were drawn from a uniform distribution that randomly varied the optimal breeding time within a ±10% range. During the climate change period, noise was drawn from a normal distribution with a mean equal to the deterministic advancement (0.1 day earlier per season) and a standard deviation of 1 day (for higher fluctuation value 1.5 and 2.5 days, please see Supplement Fig. 4). Additional simulations were also run over a longer time scale for one million seasons, to assess the stability of reproductive strategy and the consistency of our conclusions (Supplement Fig. 11). The simulation code was written in C++ (version 22) and is freely available (Zenodo repository: https://doi.org/10.5281/zenodo.21934518). Simulation data were analysed and visualised in R (version 4.2.2). This theoretical study did not require ethical approval.

