## Supplementary Figures for "Social information use promotes breeding synchrony but constrains adaptation to rapid environmental changes"

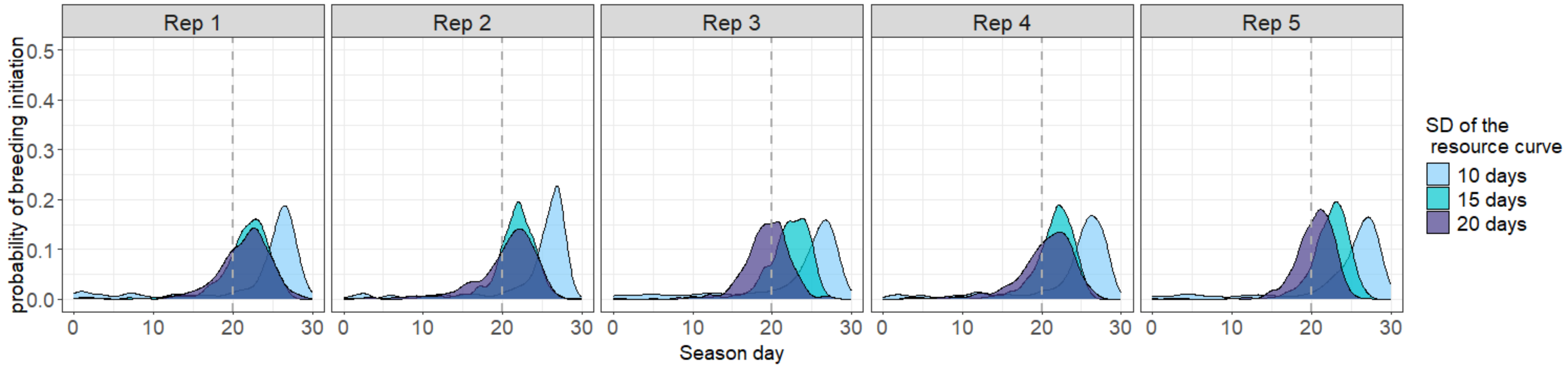


**Supplementary Figure 1**. **Breeding phenology emerged from ecological phenology with different resource variations (standard deviation in the resource curve reflecting the width of the resource peak in days; scenario i).** The standard deviation of the resource curve was set at 10 days (the same as the main results), 15 days and 20 days. Simulation outcomes of five replicates in the last season 300k are presented in panels, in which the grey dashed line indicates the optimal breeding time (day 20) of a season.

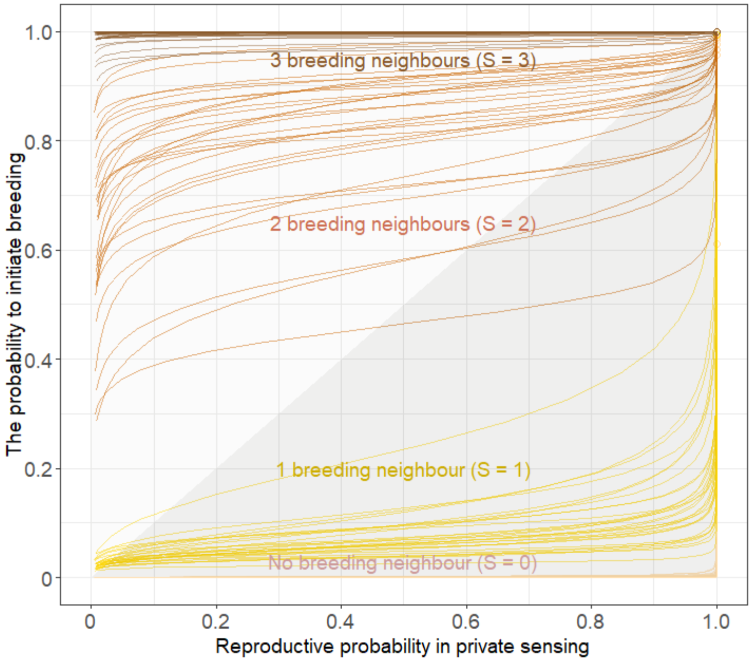


**Supplementary Figure 2. The probability to initiate breeding in the presence of social sensing is significantly enhanced relative to private sensing, when more social neighbours begin to breed.** Evolved reaction norms that reflect the probability to initiate breeding in the presence of social sensing (*y* axis) is plotted against that of private sensing (*x* axis) across 30 replicate simulations. Different colours show the number of neighbours that have begun to breed. The dark grey area implies that the probability to initiate breeding under private sensing is higher than that under social sensing, vice versa for the light grey area.

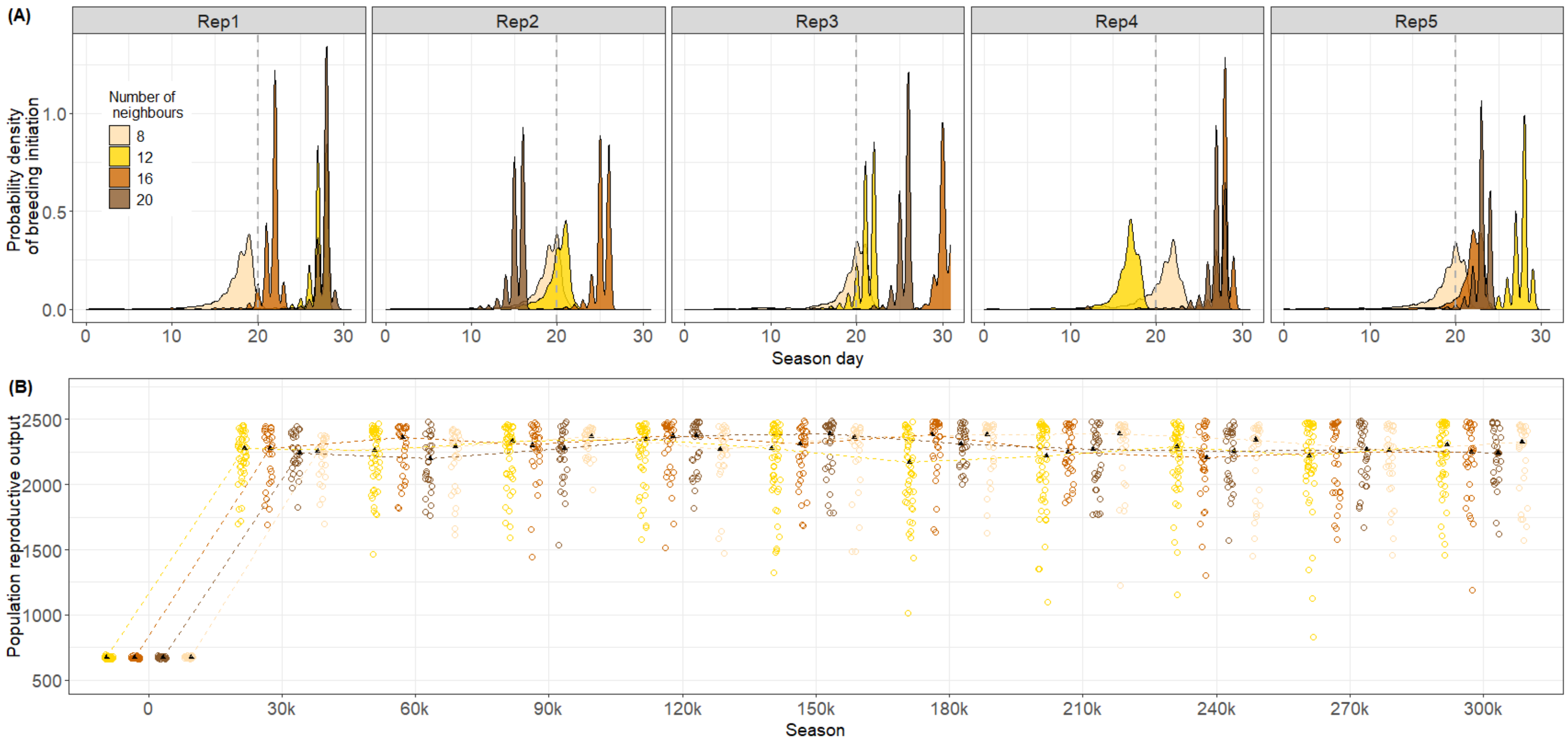


**Supplementary Figure 3. Increased neighbourhood size enhances breeding phenology peak but results in similar population reproductive output (scenario i).** Simulation outcomes when individuals are assigned 8, 12, 16, or 20 neighbours, and the corresponding ecological curve in each simulation scenario is rescaled according to the number of neighbours. Each scenario has been run for 30 replicates. (A) Breeding phenology across populations with different neighbourhood sizes at the final season (season 300k) of five random simulation replicates. Probability density at the breeding-time mode for neighbourhood sizes of 8, 12, 16 and 20, respectively: $P_8$ = 0.55±0.24, $P_{12}$ = 0.95±0.19, $P_{16}$ = 1.03±0.21, $P_{20}$ = 1.08±0.26; (B) Population reproductive output under different neighbourhood sizes. The circle present the population reproductive output from one simulation, black triangle dots show the median value across simulations, and the dashed lines link population medians to show the trends over simulation time.

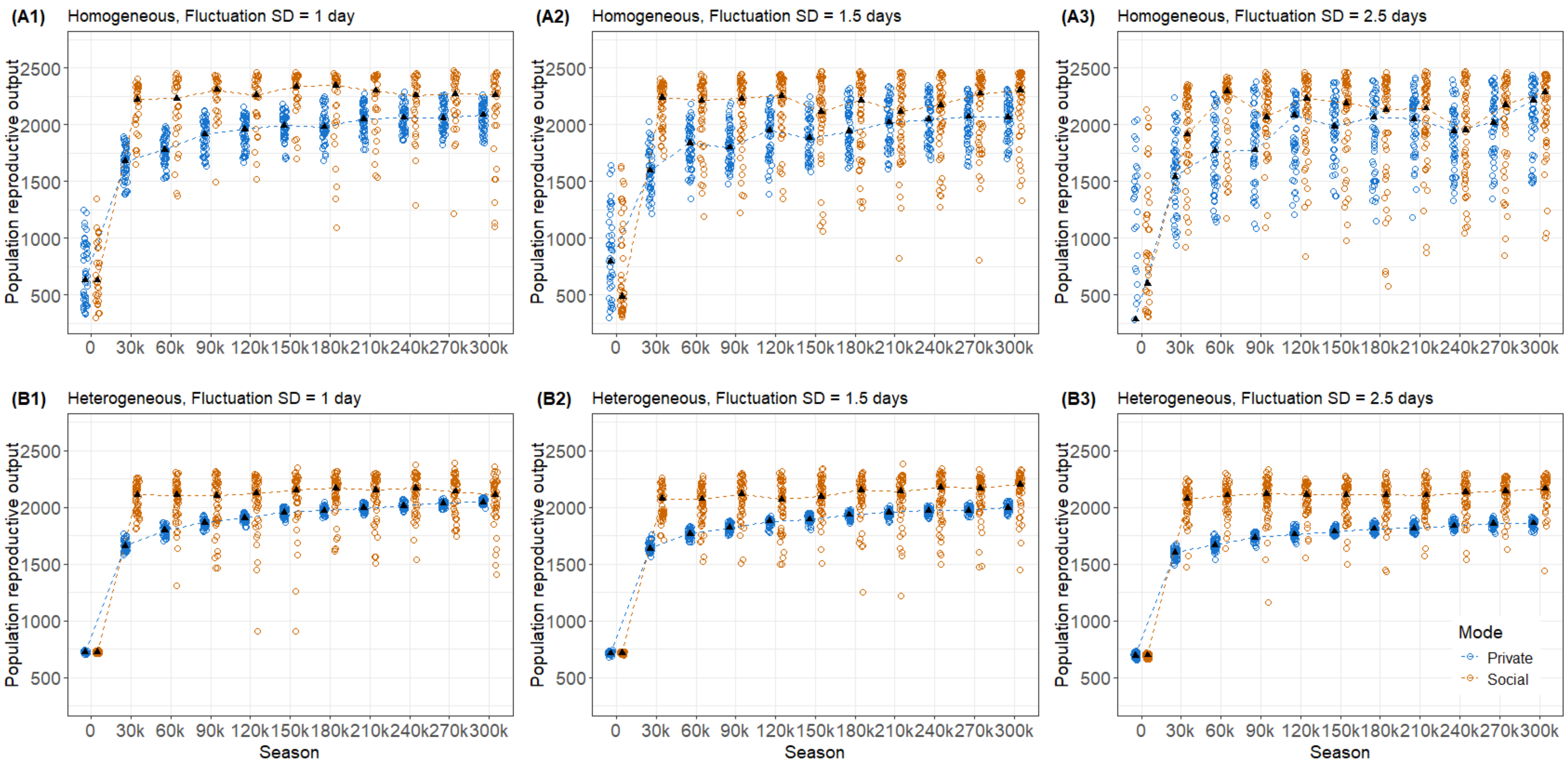


**Supplementary Figure 4. The benefits of social sensing to population performance decrease with increasing environmental noise in homogeneous landscapes, but increase in heterogeneous landscapes (Scenario ii and iv).** Comparisons of population reproductive output over the evolutionary time scale of 300K seasons between private sensing (blue dots) and social sensing (orange dots) scenarios for 50 simulation replicates each. (A) Homogeneous landscape and (B) heterogeneous landscapes with increased annual fluctuations in optimal breeding time around the average at day 20, as reflected by standard deviations of 1 day (left panels), 1.5 days (middle panels) and 2.5 days (right panels), respectively. The black dots reflect the median population reproductive output across simulations. The dashed lines linking the dots indicate the trend over environmental changes.

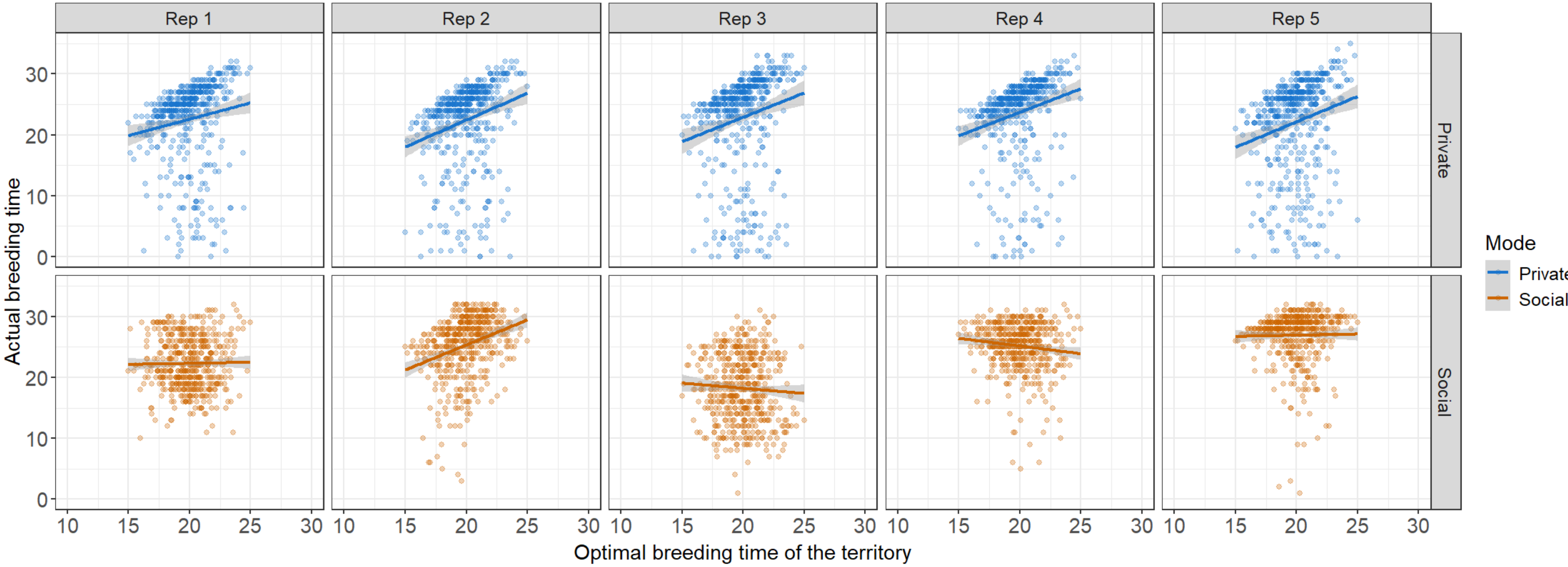


**Supplementary Figure 5. The actual breeding time is positively related to the territorial optimal breeding time in the private (blue dots), but shows no clear correlation in the social sensing (orange dots) scenario iii**. The distribution of actual breeding time over territories was plotted using outcomes from five simulation replicates. The solid lines show the trends of breeding time across territories. The grey areas indicate the 95% confidence level interval for predictions from the linear regressions.

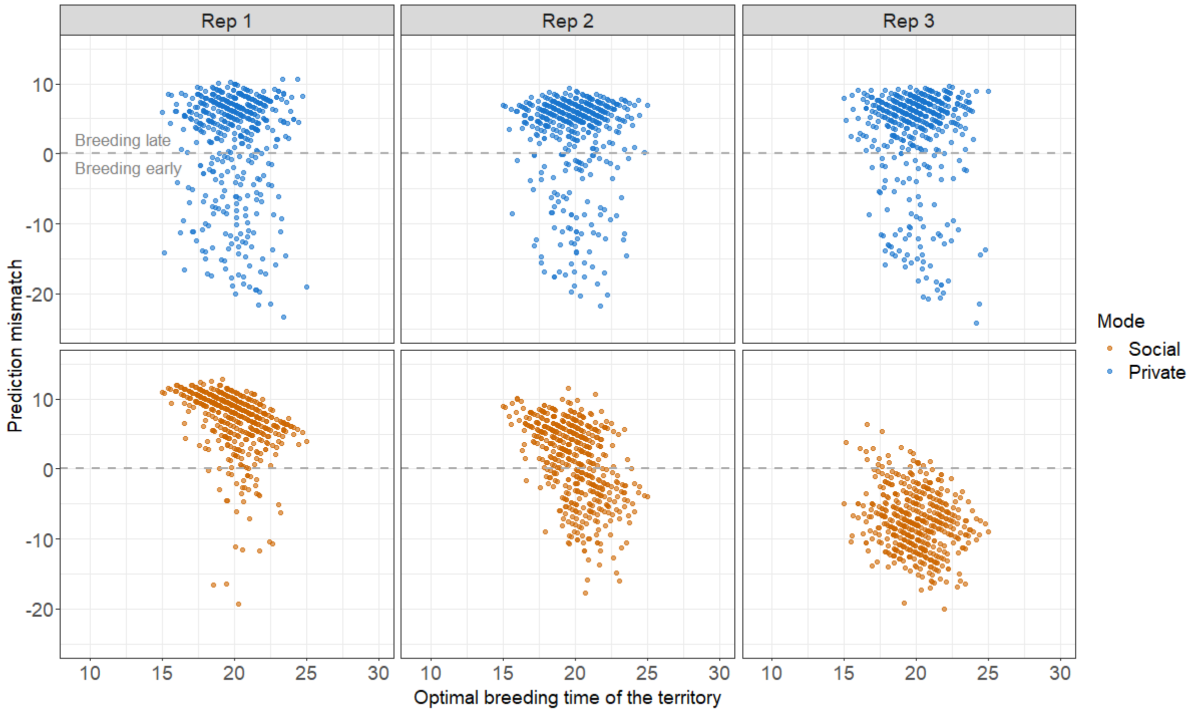


**Supplementary Figure 6. Breeding accuracy of early-resource territories leading the distribution of breeding time varied in the social sensing scenario, but not in the private sensing (scenario iii).** Three selected simulation replicates out of 30 replicates to show how a (1) late, (2) middle, and (3) early breeding prediction of breeders in early territories shape the population distribution pattern of breeding time.

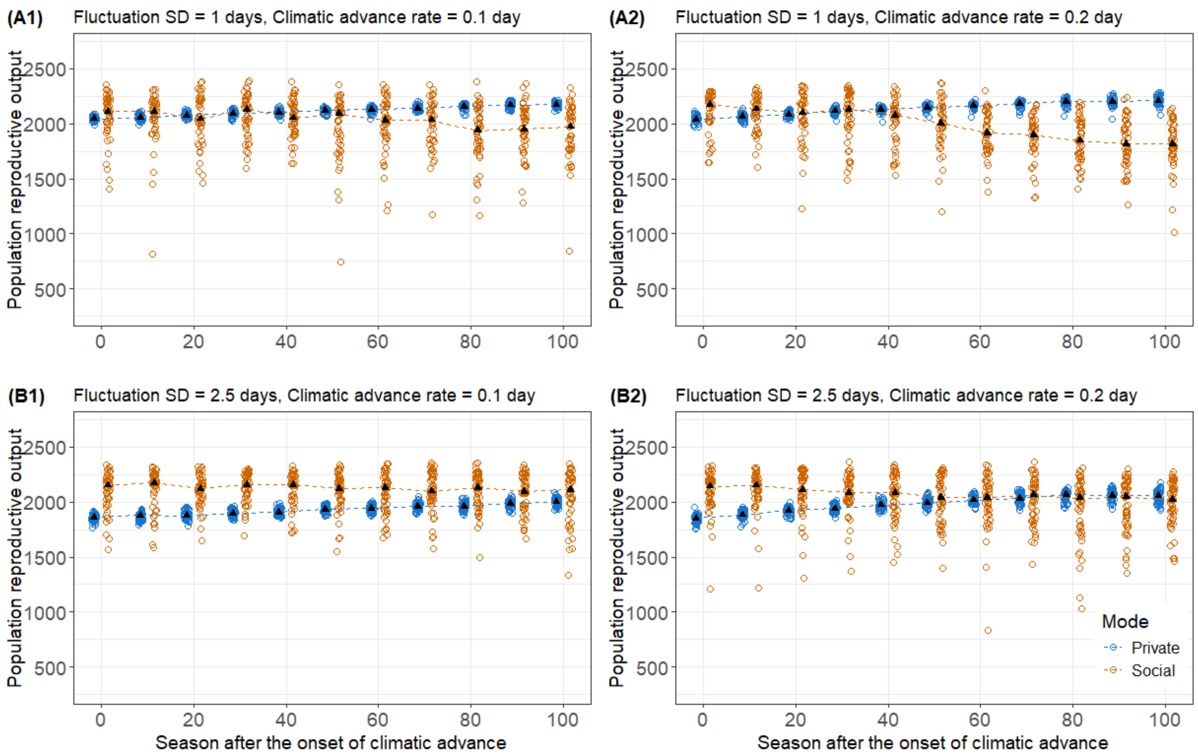


**Supplementary Figure 7. Social sensing decreases more during faster climate change, but less under larger fluctuations across seasons.** Comparison of reproductive fitness of populations breeding during different climatic advancing rates (0.1 vs 0.2 day per year) in heterogeneous and fluctuating environments (annual fluctuation SD: 1 day vs 2.5 days; scenario iv). A1 shows the same simulation outcomes as the main results in Figure 5D. The climate change phase lasts for 150 years, and both private and social sensing modes are run for 50 replicates across the four environments. The black dots refer to the median population output across simulations, and the dashed lines linking the dots indicate the trend over environmental changes.

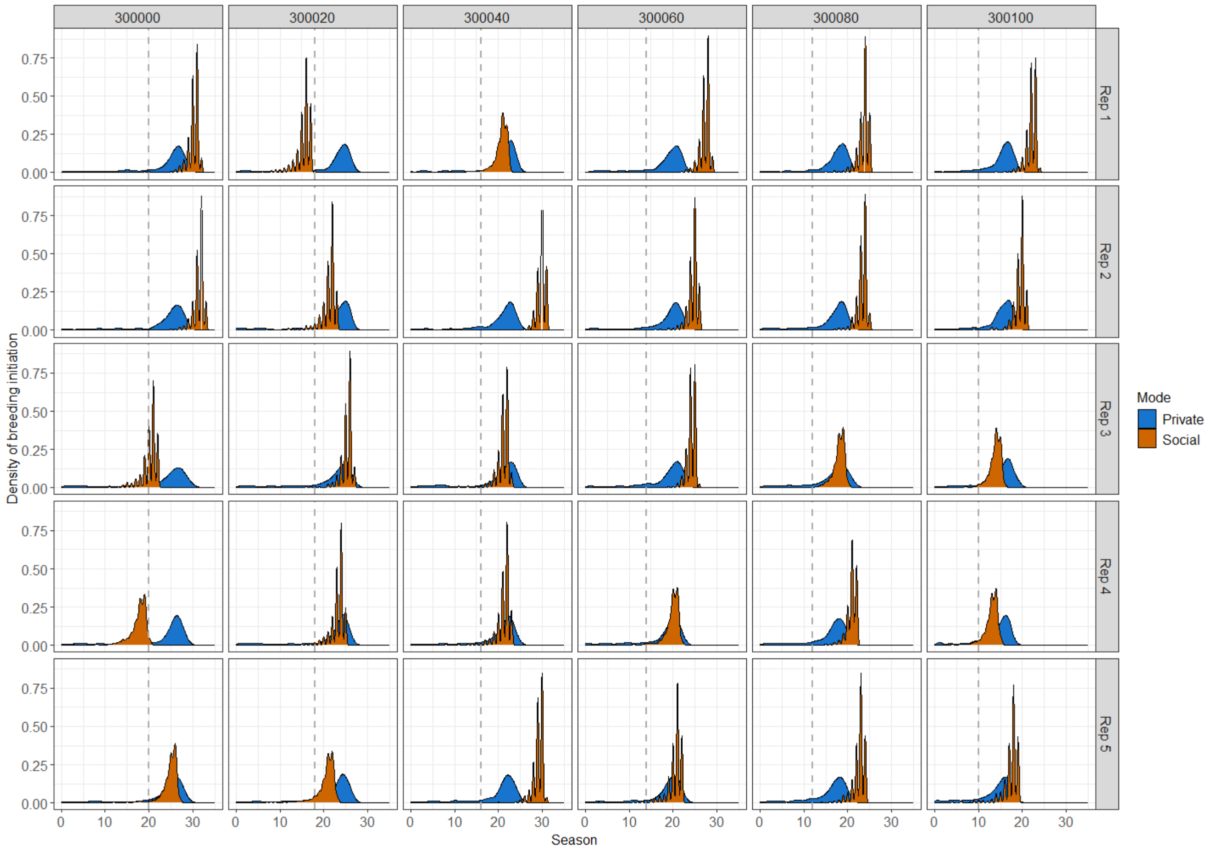


**Supplementary Figure 8. Simulation replicates of the breeding phenology in seasons during the climate change period in a homogeneous scenario (scenario i).** Outcomes of private sensing are in blue, and those of social sensing are in orange. The grey vertical dashed line shows the optimal breeding time of the season during climate change.

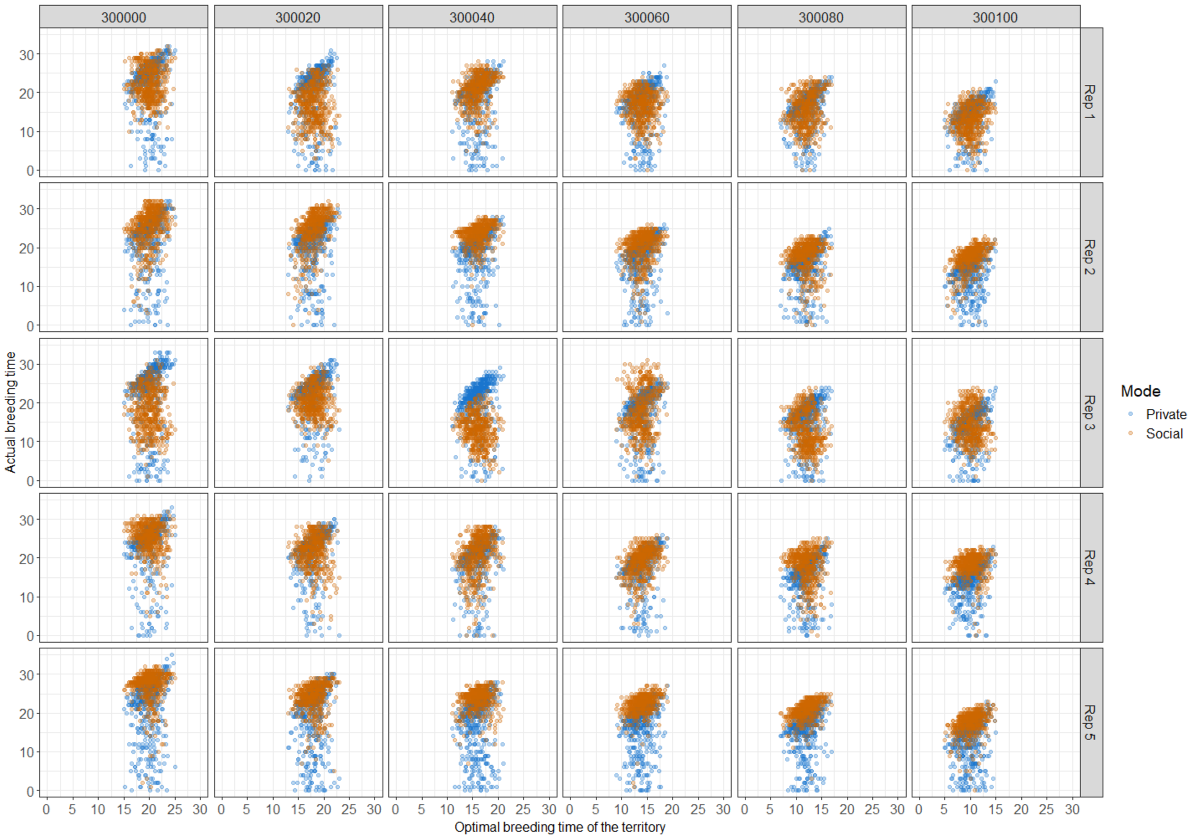


**Supplementary Figure 9. Simulation replicates of breeding time distributions in seasons during the period of climate change in the heterogeneous spatial model scenario (scenario iii).** Actual breeding time of individuals under private sensing mode is in blue, and that of social sensing mode is in orange.

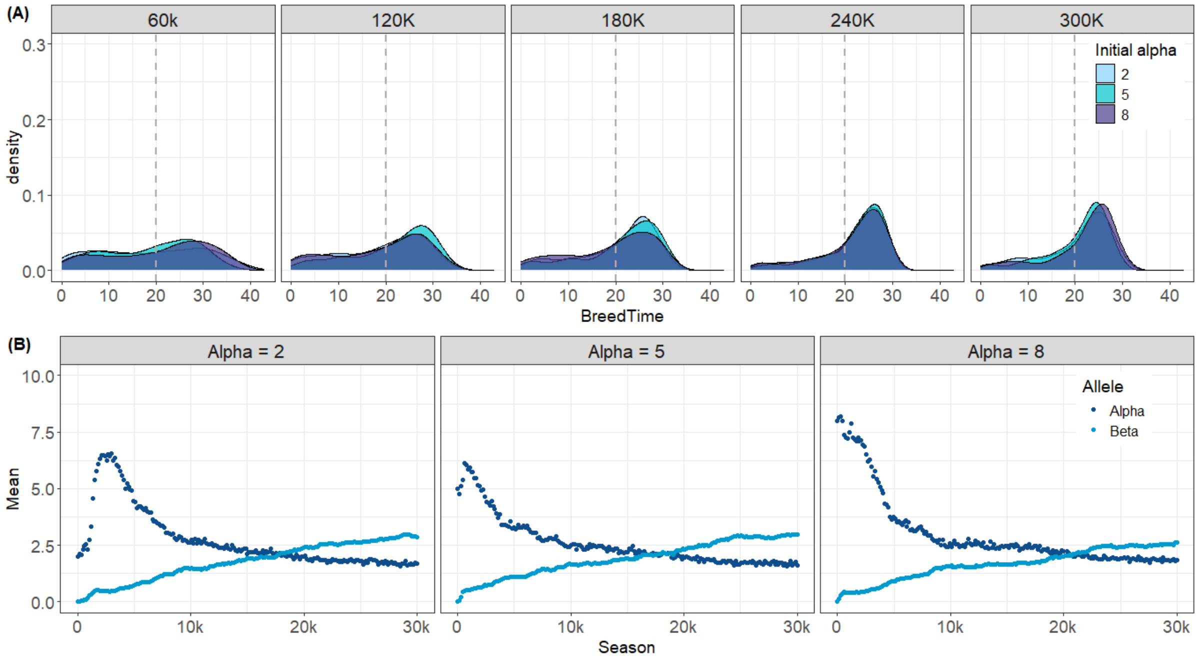


**Supplementary Figure 10. Different initial alpha values do not affect the formation of breeding phenology and allele evolution in the reproductive strategy (scenario i)**. Model initial conditions ($\alpha$ = 1, 5 or 8) are simulated 30 times each. (A) Breeding phenology that emerges over evolutionary time scales is similar across initial conditions; (B) A random simulation outcome illustrates that the evolutionary trajectories do not differ across initial conditions. The dots and error bars (very short) show population average values (mean$\pm$ SD) of $\alpha$ and $\beta$ alleles over simulation time.

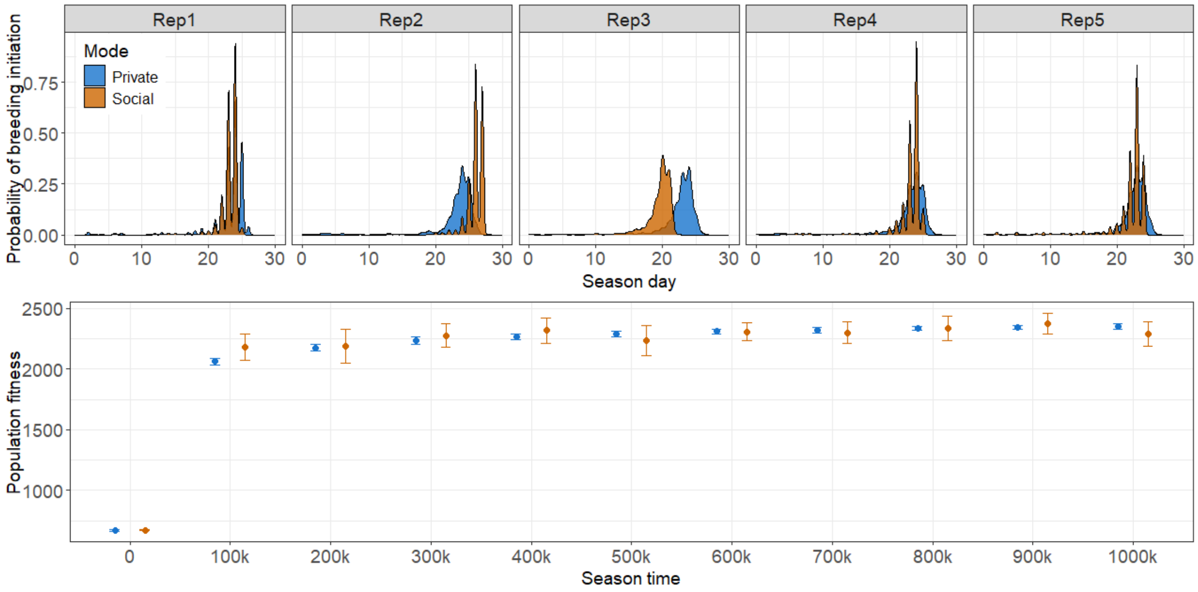


**Supplementary Figure 11. Breeding phenology under private and social sensing modes of long simulation runs for a million seasons (scenario i).** (A) The breeding phenology of five random replicates in season 1000k; (B)Population average reproductive output of 15 simulation replicates. The solid dots present the average value, and the bars indicate standard deviations.
